# A mutualistic model bacterium is lethal to non-symbiotic hosts via the type VI secretion system

**DOI:** 10.1101/2024.12.13.628426

**Authors:** Keegan E. Gaddy, Alecia N. Septer, Karen Mruk, Morgan E. Milton

## Abstract

What makes a bacterium pathogenic? Since the early days of germ theory, researchers have categorized bacteria as pathogens or non-pathogens, those that cause harm and those that do not, but this binary view is not always accurate. *Vibrio fischeri* is an exclusive mutualistic symbiont found within the light organs of Hawaiian bobtail squid. This symbiotic interaction requires *V. fischeri* to utilize a range of behaviors and produce molecules that are often associated with pathogenicity. This juxtaposition of employing “pathogenic” behaviors for a symbiotic relationship led the field to focus on how *V. fischeri* establishes a beneficial association with its host. In this study, we observe that *V. fischeri* induces mortality in zebrafish embryos and *Artemia* nauplii. Non-lethal doses of *V. fischeri* leads to zebrafish growth delays and phenotypes indicative of disease. Our data also provide evidence that the conserved type VI secretion system on chromosome I (T6SS1) plays a role in the *V. fischeri*-induced mortality of zebrafish embryos and *Artemia* nauplii. These results support the hypothesis that the *V. fischeri* T6SS1 is involved in eukaryotic cell interactions. Despite its traditional view as a beneficial symbiont, we provide evidence that *V. fischeri* is capable of harming aquatic organisms, indicating its potential to be pathogenic toward non-symbiotic hosts.

## Introduction

The *Vibrio* genus encompasses diverse marine bacteria found globally with species exhibiting free-living, symbiotic, or pathogenic lifestyles (1). As a powerful model organism for bacteria-host interactions, *Vibrio* (*Aliivibrio*) *fischeri* has been extensively studied for its mutualistic relationship with the Hawaiian bobtail squid, *Euprymna scolopes* (2–4).

Interestingly, many of the *V. fischeri* processes involved in symbiosis parallel pathogen-host interactions (2). During symbiosis establishment, *V. fischeri* has been shown to release lipopolysaccharide (LPS), peptidoglycan monomers, and small RNAs (2, 5, 6) to direct host development, form biofilm, evade immune cells, and engage in intraspecific competition via a type VI secretion (T6SS) (2). As such, *V. fischeri* challenges the conventional view of pathogenicity by employing “pathogenic” mechanisms for beneficial symbiosis. Yet a critical question remains: If *V. fischeri* is equipped with the tools of a pathogen, what prevents it from exhibiting harmful behavior?

Fish and shrimp are animals that *V. fischeri* encounter in its natural habitat as it transitions between symbiotic hosts. Zebrafish (*Danio rerio*) are an ideal model organism for studying bacterial infections and the host immune response, and have been used to explore the pathogenesis and virulence of disease causing *Vibrio* species including *Vibrio cholerae, Vibrio vulnificus*, and *Vibrio parahaemolyticus* (7). Moreover, *Artemia* nauplii have been used as an aquatic host to test pathogenicity of *V. parahaemolyticus* and *Vibrio coralliiliticus* (8, 9).

Therefore, we used zebrafish embryos and *Artemia* nauplii to address the question of potential *V. fischeri* pathogenicity. Since the T6SS is an important virulence factor for many pathogenic bacteria (10, 11), we tested the conserved T6SS on *V. fischeri* chromosome I (T6SS1) for its potential role in pathogenicity. To date, no role has been linked to the T6SS1, but it is hypothesized to be involved in eukaryotic cell interaction (12). Here, we provide evidence that directly supports this hypothesis. Our results set a new precedent by indicating there are conditions under which *V. fischeri* acts as a pathogen, expanding the field into the realm of pathogenicity.

## Results and Discussion

The impact of *V. fischeri* ES114 exposure on zebrafish embryos was tested using a bath immersion infection model. Zebrafish mortality was dose dependent and increased with the total immersion time (Fig. 1A). To determine whether embryo mortality is caused by direct or indirect effects of *V. fischeri* exposure, several control scenarios were tested. Zebrafish immersed in equivalent concentrations of *Escherichia coli* all survived (Fig. 1B), suggesting that mortality is not generally due to the presence of bacteria. We confirmed that zebrafish survival can be recovered by treating *V. fischeri* with streptomycin or heat killing prior to exposure (Fig. 1B), indicating live cells are required for mortality. Finally, to test whether mortality is caused by the accumulation of extracellular compounds, E3 media incubated with *V. fischeri* overnight was filter sterilized before embryo immersion. Survival rates were similar to untreated embryos (Fig. 1B), suggesting that *V. fischeri* is not releasing compounds into the media that are lethal to the zebrafish. These results establish that the zebrafish mortality is dependent on viable *V. fischeri* interacting with the embryos.

**Figure 1:**
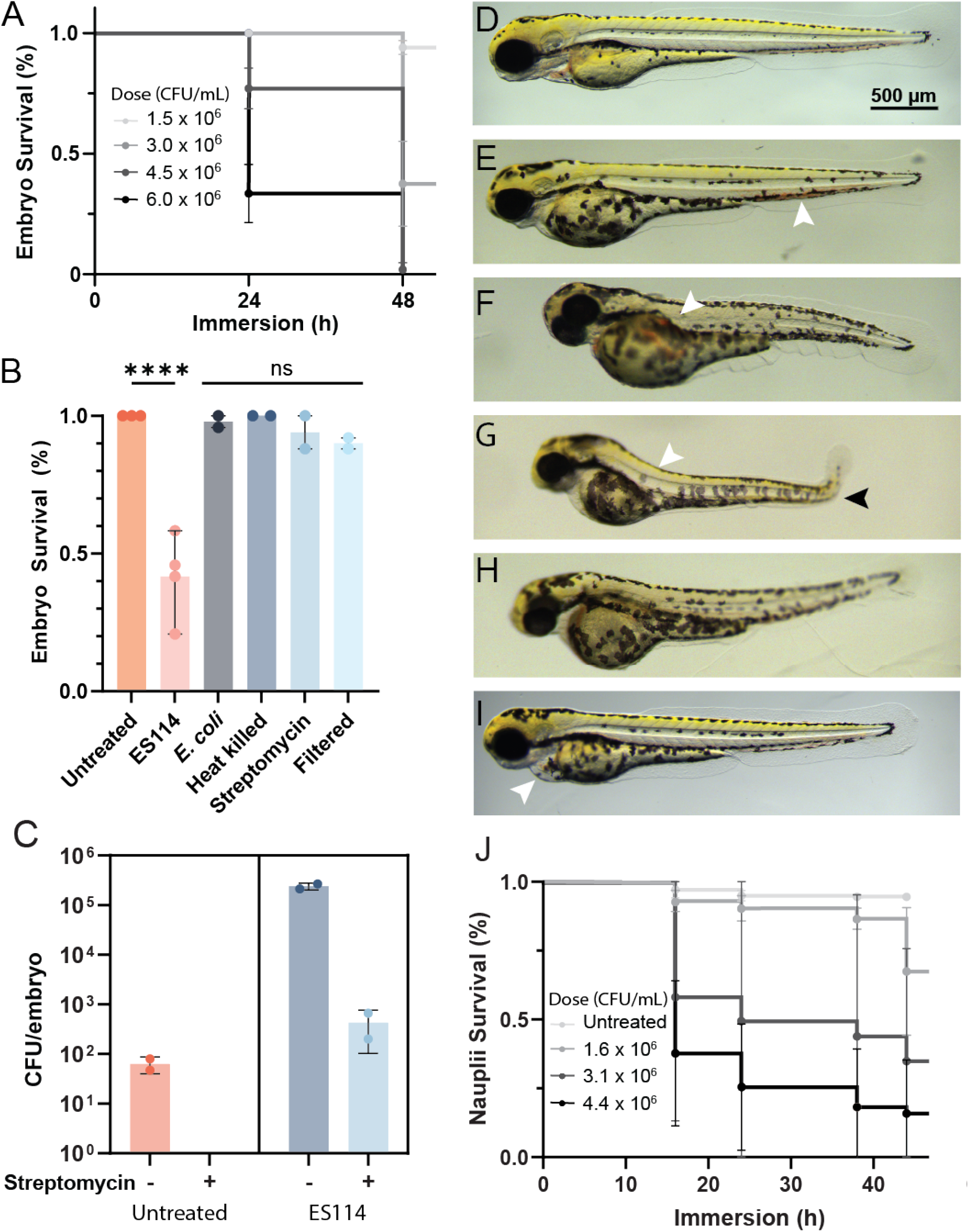
*V. fischeri* is lethal to zebrafish embryos and *Artemia* nauplii. (A) Survival curve of zebrafish embryos immersed with *V. fischeri* ES114 over 48 h. Performed with 2 independent replicates of 24 zebrafish embryos per treatment. (B) Controls used to validate dependence on viable *V. fischeri*. Each point represents independent replicates of at least 16 embryos. (C) Enumeration of bacteria following homogenization of non-sterile embryos with and without streptomycin treatment. Each point represents independent replicates of 8 embryos. (D-I) Embryos at 72 hours post fertilization (hpf) imaged following 48 h of immersion in 3.0 x 10^6^ CFU/mL *V. fischeri*. (D) Untreated control. (E) Blood pooling in the tail (arrow) and no blood flow present throughout the heart. (F) Short length with delayed yolk sac absorption, small head, necrotic tissue, and blood pooling in the duct of Cuvier (arrow). (G) Short length with severe maldevelopment including bent tail (arrow) and necrotic tissue. (H) Short length with necrotic tissue and a small head. (I) Embryo with pericardial edema. (J) Survival curve of *Artemia* nauplii immersed with *V. fischeri* ES213 over 44 h. Performed with at least 2 independent replicates with 40-100 nauplii per treatment.

Since mortality required live cells and spent supernatant was not toxic to embryos, we predicted that *V. fischeri* may be in direct contact with embryos. We evaluated *V. fischeri* associated with fish tissue after the 48 h immersion by plating and quantifying colony forming units (CFUs). Washed and homogenized embryos had approximately 2.4 x 10^5^ CFU per embryo (Fig. 1C). Embryos treated with an additional 30 min streptomycin wash prior to homogenization had approximately 4.3 x 10^2^ CFU per embryo (Fig. 1C). This indicates while many bacteria are localized to the embryo surface, a portion of *V. fischeri* may be localized within the fish tissue.

Embryos that survive *V. fischeri* exposure present with developmental defects. The most common morphological changes are shorter tail length, smaller head, necrotic tissue, and abnormal tail curvature (Fig. 1D-I). Other, less common, changes include pericardial edema (Fig. 1I). After 38 h of immersion, the most typical indicator of embryo death is slowing of the heart rate and total cessation of blood flow throughout the trunk and tail. At 48 h of immersion, most embryos displaying diminished blood flow will have no heartbeat and begin coagulation of the tail and yolk sac (Fig. 1F,H). Embryos that do not have blood flow, but still have a noticeable heartbeat, show sporadic pectoral fin movement in response to disturbance but are incapable of large movements. These progressive morphological impairments underscore the severe impact of *V. fischeri* on zebrafish embryonic development, indicating the nature of *V. fischeri* on animal development is host-specific and can range from beneficial to pathogenic (13). Exposure of another aquatic animal, *Artemia* nauplii, to *V. fischeri* ES114 revealed a similar dose dependent lethality (Fig. 1J). Taken together, the data indicate that *V. fischeri* can act as a pathogen to multiple aquatic species.

Our discovery that *V. fischeri* can indeed exhibit pathogenic behavior raises a crucial question: what virulence factors drive lethality? Given the critical role of the T6SS in pathogenicity (14), we investigated whether T6SS contributes to the pathogenic effects of *V. fischeri*. The T6SS is a molecular syringe used to deliver payloads of effectors with diverse functions. *V. fischeri* has two T6SS loci (T6SS1 and T6SS2). T6SS2 functions as an antibacterial weapon used to establish mono-colonized crypts in *E. scolopes* light organs (15). While T6SS2 is encoded by roughly half of sequenced strains, T6SS1 is conserved in all isolates, but with an unknown function (12). To determine the extent to which *V. fischeri* lethality is conserved across strains and the role of T6SS1, three symbiotic *V. fischeri* strains (ES114, ES213, and PP3) containing a deletion of the T6SS1 *icmF* gene, which encodes an essential structural protein, were compared to their wild type (WT) parent in *Artemia* nauplii (Fig 2A) and zebrafish embryo (Fig 2B) survival studies.

**Figure 2:**
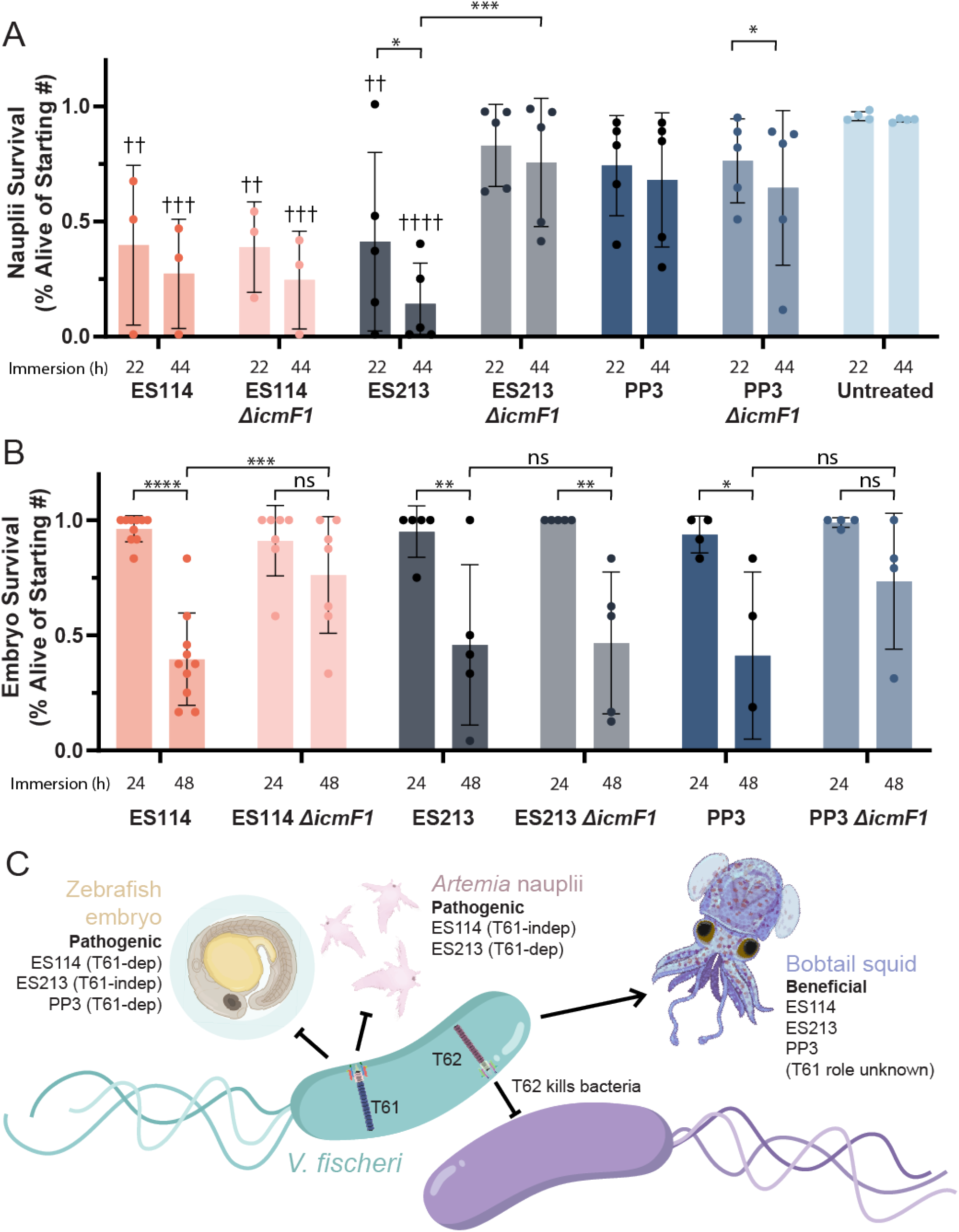
Impact of the T6SS1 on immersed animal embryo mortality. (A) Survival percentage of starting population of *Artemia* nauplii immersed in 4.6 x 10^6^ CFU/mL *V. fischeri*. Performed with at least 3 independent replicates of 40-100 nauplii per treatment. (B) Survival of zebrafish embryos immersed in 3.0 x 10^6^ CFU/mL *V. fischeri*. Untreated control embryos all survived. Performed with 4 independent replicates of 24 zebrafish embryos per treatment. The statistical significance calculated between WT, *ΔicmF1* mutants, and controls was calculated with a multiple comparisons two-way ANOVA. Asterisks show significant difference between WT and mutant pairs, daggers show significance with untreated controls, ns, no significant difference (p > 0.05), * p < 0.05, ** or †† p < 0.01, *** or ††† p < 0.001, **** or †††† p < 0.0001. (C) Model for *V. fischeri* effects on host health (pathogenic or beneficial) and the role of T6SS1, made with the help of Biorender.com.

Deletion of *icmF1* resulted in a host- and strain-specific increase in survival. Nauplii survival remained low for animals exposed to ES114 Δ*icmF1*, suggesting T6SS1 is dispensable for nauplii lethality in this strain. Nauplii showed similar levels of survival for PP3 WT or PP3 Δ*icmF1* that were not statistically different from the no-exposure control, suggesting PP3 lethality for nauplii is low. However, ES213 WT exposure resulted in low nauplii survival that was significantly higher when *icmF1* was deleted, suggesting a role for T6SS1 in ES213 lethality against *Artemia*.

Interestingly, the strain-specific lethality phenotypes were reversed for zebrafish embryos (Fig 2B). ES213 Δ*icmF1* retained lethality, similar to the WT parent, suggesting T6SS1 is dispensable for this strain. However, embryos exposed to the *icmF1* mutants for both ES114 and PP3 showed higher survival rates when compared to their WT parents. Although the difference between zebrafish survival for PP3 WT and PP3 Δ*icmF1* was not statistically significant, the effect size is consistent, and the *icmF1* mutant exposed embryos reached higher levels of survivability, compared to WT, for each of the four independent trials. These results suggest that, under our experimental conditions, the T6SS1 is involved in pathogenesis and supports the hypothesis that T6SS1 mediates interactions with eukaryotic hosts.

In summary, *V. fischeri’s* pathogenic behavior towards non-symbiotic hosts expands its utility as a model organism beyond that of beneficial symbiosis (Fig 2C). Future work will employ this multi-strain and multi-host model system to explore mechanistic connections between beneficial and harmful infection.

## Methods

See SI Appendix for experimental details.

## Acknowledgments

We thank Thomas Rynes for excellent fish care, Dr. Karen Visick for her generous gift of ES114 and thoughtful discussions, and AJ Milton for editing. This work was supported by NIAID (K22AI170662 to M.E.M), NIGMS (R35 GM137886 to A.N.S. and R21GM143565 to K.M.), and the Brody School of Medicine (to M.E.M. and K.M.).

## Author contributions

M.E.M. conceived research; all authors designed research; K.E.G. performed research; A.N.S. generated mutants and provided bacterial strains; K.M. provided zebrafish embryos and brine shrimp; all authors analyzed data; and all authors wrote the paper.

