## Extended Methods for "A mutualistic model bacterium is lethal to non-symbiotic hosts via the type VI secretion system"

*Zebrafish husbandry*

Zebrafish [WT AB (ZL1)] were purchased from the Zebrafish International Resource Center (ZIRC). Adults were maintained at 28.5 °C in a recirculating system (Iwaki Aquatics) on a 14:10 h light:dark cycle and fed in the morning with brine shrimp (EZ-Egg, Brine Shrimp Direct) and in the afternoon with Zeigler's Adult Zebrafish Diet (Pentair Aquatic Habitats). Embryos were obtained through natural matings and cultured at 28–30 °C in E3 medium. The National Institutes of Health Office of Laboratory Animal Welfare (OLAW) is responsible for the administration of the US Public Health Service Policy on Humane Care and Use of Laboratory Animals. Compliance with the standards of the Public Health Service Policy is a term and condition of NIH Grants Policy Statement. East Carolina University's Animal Welfare Assurance number is A3469-01. The IACUC committee at East Carolina University, Greenville, NC, USA approved all animal procedures (AUP#W262).

*Brine shrimp husbandry*

A hatchery (Pentair Aquatic Habitats) containing 75 g of sea salt (Instant Ocean) and 2 L of RO water was prepared. Twenty-four hours before the experiment, 1 teaspoon of EZ-Egg brine shrimp (Brine Shrimp Direct, Utah) was added to the hatchery and kept with continuous agitation at 28.5 °C. The next day, shrimp were collected for experimentation.

*Bacterial strain construction*

For construction of *V. fischeri icmF_1* deletion mutants (Δ*icmF_1*) in strains ES114, PP3, and ES213, the upstream and downstream regions were PCR amplified using primers AS1067/AS1069 and AS1068/AS1070, respectively, using gDNA from the parent strain. The upstream and downstream fragments were combined using SOE PCR and primers AS1067 and AS1068. The resulting product was cloned into pCR-BluntII-topo vector using a Zero Blunt PCR Cloning Kit from Thermo Fisher and fused to pEVS118 at the KpnI site. The Δ*icmF_1* mutations on these plasmids were moved into *V. fischeri* strains by triparental mating using the conjugative helper strain pEVS104. Single recombinants were verified by antibiotic resistance and double recombinants were verified by loss of antibiotic resistance and PCR verification of deletion allele.

Genotypes of plasmids were all confirmed with PCR and Sanger Sequencing. *V. fischeri* mutant construction was verified with antibiotic resistance/sensitivity and PCR validation. Details on strain genotypes, source, plasmids, and primers are in Table S1.

*Preparation of bacterial cultures*

Cultures of *V. fischeri* were grown in 100 mL unbuffered LBS broth overnight (16-18 h) at 28 °C with shaking. Overnight cultures were subcultured in 100 mL fresh unbuffered LBS broth and grown at 28 °C until reaching an $OD_{600}$ of 1.0. To collect bacteria for immersion assays, 10 mL of the subculture was spun down, washed once with E3 media, and resuspended in an equivalent volume of E3.

*Infection of zebrafish embryos*

Twenty four hours post fertilization (hpf), embryos were placed individually into sterile clear 96-well plates and immersed in the desired concentration of bacteria in E3 media to a final volume of 150 µL. For dose response experiments, washed *V. fischeri* ES114 was diluted in E3 to the final desired $OD_{600}$. For the five control experiments, embryos were placed in either sterile E3 media, media with heat-killed *V. fischeri* ES114, media with *V. fischeri* treated with 0.25 mg/mL streptomycin, E3 media exposed to *V. fischeri* overnight and then filtered through a 0.22 µm PVDF filter, or BL21 *E. coli*. When testing the impacts of different strains and mutants, washed *V. fischeri* was diluted to a final $OD_{600}$ of 0.4 (approximately 3 x $10^6$ CFU/mL). For all conditions, mortality and other growth defects were observed at 24 and 48 h post-immersion.

For conditions using heat killed bacteria, 1 mL volumes of washed *V. fischeri* cultures in E3 were placed on a heat block at 80 °C for 20 minutes before allowing to cool to room temperature. The heat-treated media was plated and incubated overnight at 28 °C to ensure cell death. For filtered E3 media, *V. fischeri* were grown overnight in unbuffered LBS at 28 °C. Cells were then washed and subculture into E3 and allowed to incubate overnight at 28 °C, shaking. The media was then passed through a 0.22 µm PVDF filter before exposing the embryos. For *E. coli* immersion, the *E. coli* was grown in LB to an OD of 1.0, pelleted, washed, and resuspended in E3 media to replicate *V. fischeri* treatments.

*Zebrafish homogenization and bacterial load*

Colony-forming units (CFUs) were determined by plating three 5 µL spots of serial diluted sample on unbuffered LBS agar and incubating overnight at 28 °C. To determine CFUs from immersed embryos, six embryos were washed twice in fresh E3 media and placed in a single microcentrifuge tube with glass beads and 200 µL PBS. A subset of embryos were exposed to 0.25 mg/mL streptomycin for 30 min and washed again in fresh E3 media before being transferred in microcentrifuge tubes. To macerate embryos, the microcentrifuge tubes were placed on a vortex mixer until homogeneous. The homogenate was serial diluted with PBS and spot plated on unbuffered LBS agar to determine CFUs.

*Infection of Artemia*

Hatched *Artemia* nauplii were transferred into sterile 6-well plates containing 4 mL of Instant Ocean with 30-100 nauplii per well. Nauplii were challenged with *V. fischeri* in the same manner as described above for zebrafish. Plates were incubated at 28 °C. Survival was determined at the indicated time points post infection. A nauplii that did not move within 10 s was defined as nonviable. Percent survival was calculated for each time point as the surviving nauplii out of the total.

*Microscopy*

After experimentation, zebrafish embryos were screened using a Revolve Echo equipped with Olympus 4x Plan Fluorite Objective lens. Live embryos at 72 hpf were mounted on glass depression slides using 1.5% LMP agarose. Embryos were imaged on a Leica M165 equipped with a K7 color CMOS camera. Images were saved as TIFFs and processed in Photoshop (Adobe). Adjustments were limited to levels and cropping.

*Statistical analysis*

For all zebrafish experiments, at least two breeding tanks, each containing 2-3 males and 3-5 females from separate stocks, were set up to generate embryos. Embryos from each tank were randomly distributed across tested conditions. Unfertilized eggs and developmentally abnormal embryos were removed prior to treatment or imaging. Power analysis was conducted to determine the number of embryos needed in each condition to observe a 50% size effect with 80% power. Data were analyzed using GraphPad Prism (version 10.2.3). Survival curves of dose-dependent effects were calculated using Kaplan-Meier analysis with significance determined by the Mantel-Cox test. Unpaired t-tests were used for comparison of mortality between *V. fischeri* strains. A *P* value of <0.05 was used for all statistical significance.

**Table S1. *V. fischeri* strains, plasmids, and primers used in this study.**

| Strains or Plasmids | Relevant characteristics | Source or Ref. |
|---|---|---|
| ***E. coli*** | | |
| DH5α | F'/*endA1 hsdR17 glnV44 thi-1 recA1 gyrA relA1* Δ(*lacIZYA-argF*)*U169deoR*(f80*dlacI*Δ(*lacZ*)*M15*) | Hanahan 1983 (1) |
| DH5αλ pir | λ*pir* derivative of DH5α | Dunn et al. 2005 (2) |
| CC118λpir | Δ(*ara-leu*) *araD* Δ*lac74 galE galK phoA20 thi-1 rpsE rpsB argE*(Am) *recA* λ*pir* | Herrero et al. 1990 (3) |
| ***V. fischeri***[a] | | |
| ES114 | *Euprymna scolopes* light organ isolate, Kaneohe Bay, HI | Boettcher & Ruby 1990 (4) |
| ES213 | *Euprymna scolopes* light organ isolate, Maunalua Bay, HI | Boettcher & Ruby 1994 (5) |
| PP3 | Planktonic isolate, Kaneohe Bay, HI | Lee & Ruby 1992 (6) |
| ANS2034 | PP3 Δ*icmF_1* | This study |
| ANS2040 | ES114 Δ*icmF_1* | This study |
| ANS2046 | ES114 Δ*icmF_1* pVSV208 | This study |
| ANS2093 | ES213 Δ*icmF_1* | This study |
| **Plasmids** | | |
| pCR-BluntII-topo | *oriV*$_{ColE1}$, Kn$^R$ | Thermo Fisher |
| pAS2000 | ES114 Δ*icmF_1* allele in topo vector; *oriV*$_{ColE1}$, Kn$^R$ | This study |
| pAS2006 | PP3 Δ*icmF_1* allele in topo vector; *oriV*$_{ColE1}$, Kn$^R$ | This study |
| pAS2015 | ES114 Δ*icmF_1* allele (pAS2000 fused to pEVS118) *oriV*$_{R6K\gamma}$, *oriV*$_{ColE1}$, *oriT*, Cm$^R$, Kn$^R$ | This study |
| pAS2018 | PP3 Δ*icmF_1* allele (pAS2006 fused to pEVS118) *oriV*$_{R6K\gamma}$, *oriV*$_{ColE1}$, *oriT*, Cm$^R$, Kn$^R$ | This study |
| pEVS104 | conjugative helper, *oriV*$_{R6K\gamma}$, *oriT*, Kn$^R$ | Stabb & Ruby, 2002 (7) |
| pEVS118 | *oriV*$_{R6K\gamma}$, *oriT,* Cm$^R$ | Dunn et al., 2005 (2) |
| pEVS122 | *oriV*$_{R6K\gamma}$, *oriT,* Erm$^R$ | Dunn et al., 2005 (2) |
| pVSV208 | *dsRed+, oriV*$_{R6K\gamma}$, *oriV*$_{pES213}$, *oriT*, Cm$^R$ | Dunn et al., 2006 (8) |
| **Oligos**[a] | | |
| AS1067 | GAAGCCACCTTTATTCTCGCGGC | This study |
| AS1068 | TTCACCTTTAATAACCGGCATAATAAGAAAACC | This study |
| AS1069 | AATGGGCCGTTTGAAATTGTTGGCTCATCCGTAAAATCC TATAAATCAATTTGCTG | This study |
| AS1070 | CAGCAAATTGATTTATAGGATTTTACGGATGAGCCAACA ATTTCAAACGGCCCATT | This study |

[a]Restriction sites are underlined.
